# Rate-distortion theory of neural coding and its implications for working memory

**DOI:** 10.1101/2022.02.28.482269

**Authors:** Anthony M.V. Jakob, Samuel J. Gershman

## Abstract

Rate-distortion theory provides a powerful framework for understanding the nature of human memory by formalizing the relationship between information rate (the average number of bits per stimulus transmitted across the memory channel) and distortion (the cost of memory errors). Here we show how this abstract computational-level framework can be realized by a model of neural population coding. The model reproduces key regularities of visual working memory, including some that were not previously explained by population coding models. We verify a novel prediction of the model by reanalyzing recordings of monkey prefrontal neurons during an oculomotor delayed response task.

## Introduction

All memory systems are capacity-limited in the sense that a finite amount of information about the past can be stored and retrieved without error. Most digital storage systems are designed to work without error. Memory in the brain, by contrast, is error-prone. In the domain of working memory, these errors follow well-behaved functions of set size, variability, attention, among other factors. An important insight into the nature of such regularities was the recognition that they may emerge from maximization of memory performance subject to a capacity limit or encoding cost [1, 2, 3, 4, 5, 6, 7].

Rate-distortion theory [8] provides a general formalization of the memory optimization problem (reviewed in [9]). The costs of memory errors are specified by a *distortion function*; the capacity of memory is specified by an upper bound on the mutual information between the inputs (memoranda) and outputs (reconstructions) of the memory system. Systems with higher capacity can achieve lower expected distortion, tracing out an optimal trade-off curve in the rate-distortion plane. The hypothesis that human memory operates near the optimal trade-off curve allows one to deduce several known regularities of working memory errors, some of which we describe below. Past work has studied rate-distortion trade-offs in human memory [1, 2, 10], as well as in other domains such as category learning [4], perceptual identification [11], visual search [5], linguistic communication [12], and decision making [13, 14].

Our goal is to show how the abstract rate-distortion framework can be realized in a neural circuit using population coding. As exemplified by the work of Bays and his colleagues, population coding offers a systematic account of working memory performance [15, 16, 17, 18, 19, 20, 21], according to which errors arise from the readout of a noisy spiking population that encodes memoranda. We show that a modified version of the population coding model implements the celebrated Blahut-Arimoto algorithm for rate-distortion optimization [22, 23]. The modified version can explain a number of phenomena that were puzzling under previous population coding accounts, such as *serial dependence* (the influence of previous trials on performance [24]).

The Blahut-Arimoto algorithm is parametrized by a coefficient that specifies the trade-off between rate and distortion. In our circuit implementation, this coefficient controls the precision of the population code. We derive a homeostatic learning rule that adapts the coefficient to maintain performance at the capacity limit. This learning rule explains the dependence of memory performance on the intertrial and retention intervals [25, 26, 27]. It also makes the prediction that performance should adapt across trials to maintain a set point close to the channel capacity. We confirm these performance adjustments empirically. Finally, we show that variations in performance track changes in neural gain, consistent with our theory.

## Results

### The channel design problem

We begin with an abstract characterization of the channel design problem, before specializing it to the case of neural population coding. A communcation channel (Figure 1A) is a probabilistic mapping, 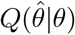, from input *θ* to a reconstruction 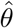. The input and output spaces are assumed to be discrete in our treatment (for continuous variables like color and orientation, we use discretization into a finite number of bins; see also [2]). We also assume that there is some capacity limit *C* on the amount of information that this channel can communicate about *θ*, as quantified by the mutual information 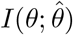 between *θ* and the stimulus estimate 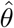 decoded from the population activity. We will refer to 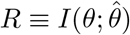 as the channel’s *information rate*. To derive the optimal channel design, we also need to specify what *distortion function* 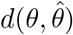 the channel is optimizing—i.e., how errors are quantified. Details on our choice of distortion function can be found below.

**Figure 1:**
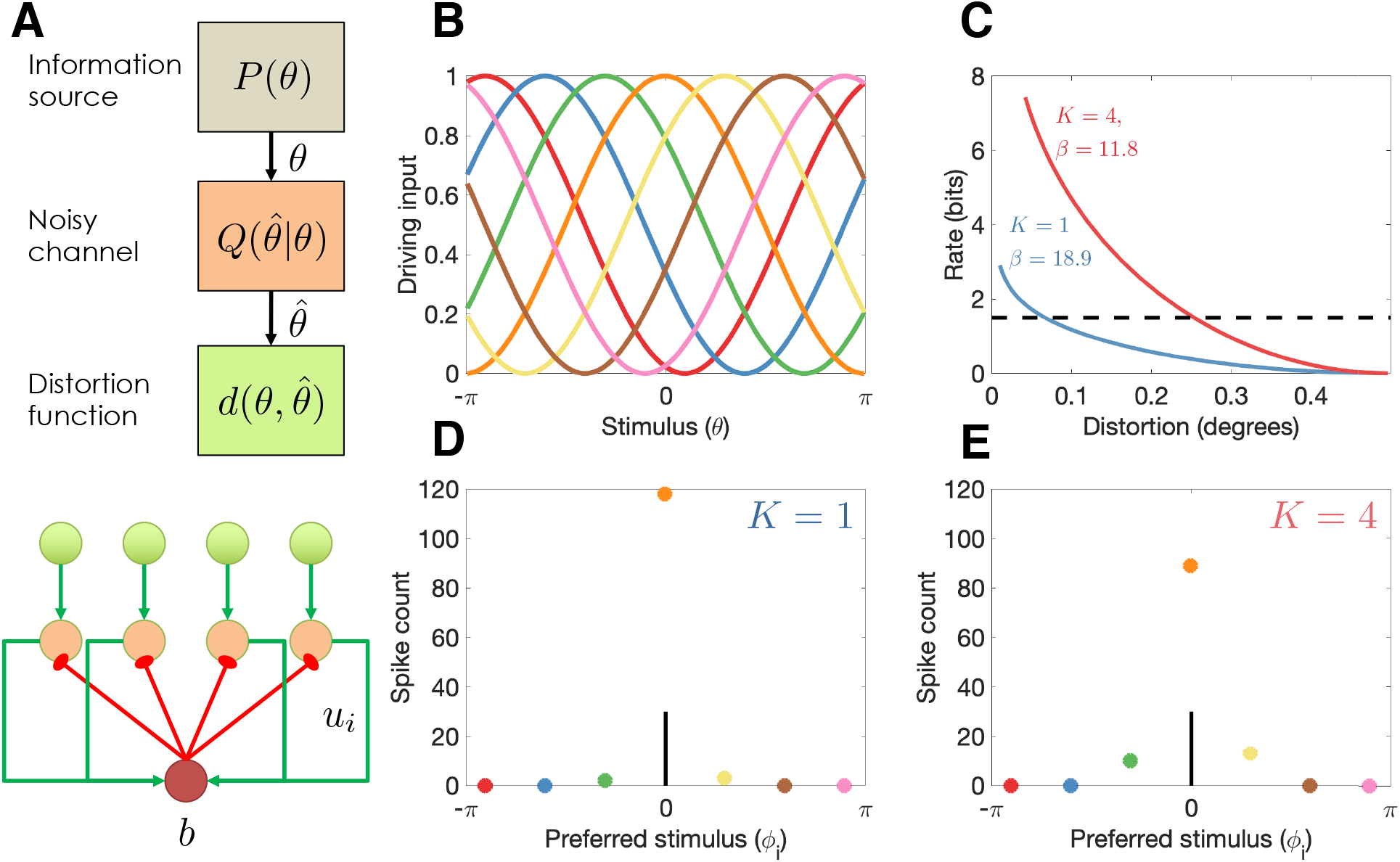
Model illustration. (A) Top: Abstract characterization of a communication channel. A stimulus *θ* is sampled from an information source *P* (*θ*) and passed through a noisy communication channel 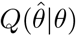, which outputs a stimulus reconstruction 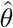. The reconstruction error is quantified by a distortion function, 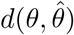. Bottom: Circuit architecture implementing the communication channel. Input neurons encoding the negative distortion function provide the driving input to output neurons with membrane potential *u*_*i*_ and global feedback inhibition *b*. Each circuit codes a single stimulus at a fixed retinotopic location. When multiple stimuli are presented, the circuits operate in parallel, interacting only through a common gain parameter, *β*. (B) Tuning curves of input neurons encoding the negative cosine distortion function over a circular stimulus space. (C) Rate-distortion curves for two different set sizes (*K* = 1 and *K* = 4). The optimal gain parameter *β* is shown for each curve, corresponding to the point at which each curve intersects the channel capacity (horizontal dashed line). Expected distortion decreases with the information rate of the channel, but the channel capacity imposes a lower bound on expected distortion. (D) Example spike counts for output neurons in response to a stimulus (*θ* = 0, vertical line). The output neurons are color-coded by their corresponding input neuron (arranged horizontally by their preferred stimulus, *ϕ*_*i*_ for neuron *i*; full tuning curves are shown in panel B). When only a single stimulus is presented (*K* = 1), the gain is high and the output neurons report the true stimulus with high precision. (E). When multiple stimuli are presented (*K* = 4), the gain is lower and the output has reduced precision (i.e., sometimes the wrong output neuron fires).

With these elements in hand, we can define the channel design problem as finding the channel *Q*^∗^ that minimizes expected distortion 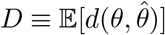 subject to the constraint that the information rate *R* cannot exceed the capacity limit *C*:

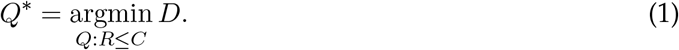

For computational convenience, we can equivalently formulate this problem using a Lagrangian:

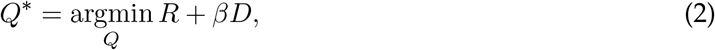

where *β* is a Lagrange multiplier equal to the slope of the rate-distortion function at the capacity limit:

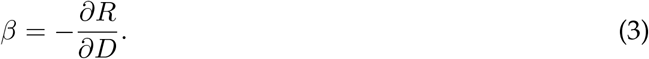

The rate-distortion function is monotonically decreasing and convex; hence the value of *β* is always positive.

Using the Lagrangian formulation, one can show that the optimal channel for a discrete stimulus takes the following form:

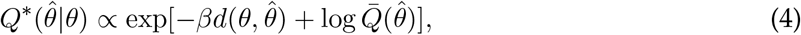

where the marginal probability 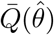 is defined by:

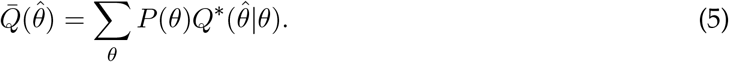

These two equations are coupled. One can obtain the optimal channel by initializing them to uniform distributions and iterating them until convergence. This is known as the Blahut-Arimoto algorithm [22, 23].

For a channel with a fixed capacity *C* but variable *D* across contexts, the Lagrange multiplier *β* will need to be adjusted for each context so that *R* = *C*. We can accomplish this by computing *R* for a range of *β* values and choosing the value that gets closest to the constraint *C* (later we will propose a more biologically plausible algorithm). Because the rate-distortion function is monotonically decreasing and convex, *β* will always be a decreasing function of *D*. Intuitively, *β* characterizes the sensitivity of the channel to the stimulus. When stimulus sensitivity is lower, the information rate is lower and hence the expected distortion is higher.

In general, we will be interested in communicating a collection of *K* stimuli, *θ* = {*θ*_1_, …, *θ*_*K*_}, with associated probing probabilities *π* ={ *π*_1_, …, *π*_*K*_ }, where *π*_*k*_ is the probability that stimulus *k* will be probed [3]. The resulting distortion function is obtained by marginalizing over the probe stimulus:

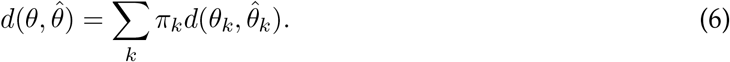

### Optimal population coding

We now consider how to realize the optimal channel with a population of spiking neurons, each tuned to a particular stimulus (Figure 1A). The firing rate of neuron *i* is determined by a simple Spike Response Model [28] in which the membrane potential is the difference between the excitatory input, *u*_*i*_, and the inhibitory input, *b*, which we model as common across neurons (to keep notation simple, we will suppress the time index for all variables). Spiking is generated by a Poisson process, with firing rate modeled as an exponential function of the membrane potential [29]:

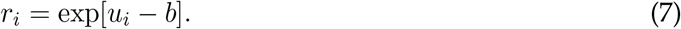

We assume that inhibition is strong enough such that only one neuron in the population will be active within an infinitesimally small time window. In particular, we assume that inhibition is given by *b* = log∑_*i*_ exp[*u*_*i*_], in which case the firing rate is driven by the excitatory input with divisive normalization [30]:

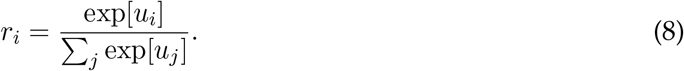

Note that allowing the feedback inhibition to be a function of the membrane potential *b* is biologically unrealistic, since the interneurons driving the inhibition do not have access to the membrane potential of other neurons. However, for our purposes it suffices to assume that the inhibitory population can approximate this function based on the spiking output of the excitatory neurons.

The resulting population dynamics is a form of “winner-take-all” circuit [31]. If each neuron has a preferred stimulus *ϕ*_*i*_, then the winner can be understood as the momentary channel output, 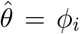 whenever neuron *i* spikes (denoted *z*_*i*_ = 1). The probability that neuron *i* is the winner within a given infinitesimal time window is:

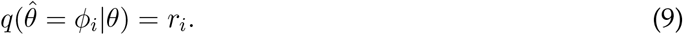

Importantly, Equation 9 has the same functional form as Equation 4, and the two are equivalent if the excitatory input is given by:

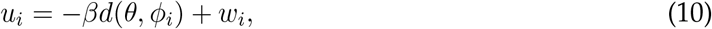

where

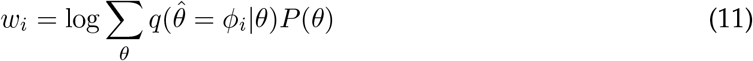

is the log marginal probability of neuron *i* being selected as the winner. We can see from this expression that the first term in Equation 10 corresponds to the neuron’s stimulus-driven excitatory input and the second term corresponds to the neuron’s excitability. The Lagrange multiplier *β* plays the role of a gain modulation factor.

The excitability term can be learned through a form of intrinsic plasticity [31], using the following spike-triggered update rule:

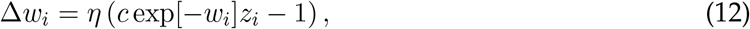

where *η* is a learning rate and *c* a gain parameter. After a spike (*z*_*i*_ = 1), the excitability is increased proportionally to the inverse exponential of current excitability. In the absence of a spike, the excitability is decreased by a constant. This learning rule is broadly in agreement with experimental studies [32, 33].

We now address how to optimize *β*. We want the circuit to operate at the set point *R* = *C*, where the channel capacity *C* is understood as some fixed property of the circuit, whereas the information rate *R* can vary based on the parameters and input distribution, but cannot persistently exceed *C*. Assuming the total firing rate of the population is approximately constant across time, we can express the information rate as follows:

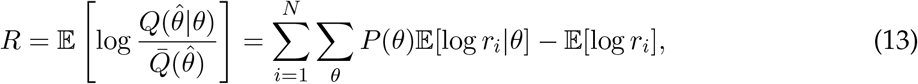

where *N* is the number of neurons. This expression reveals that channel capacity corresponds to a constraint on stimulus-driven deviations in firing rate from the marginal firing rate. When the stimulus-driven firing rate is persistently greater than the marginal firing rate, the population may incur an unsustainably large metabolic cost [34, 35]. When the stimulus-driven firing rate is lower than the marginal firing rate, the population is underutilizing its information transmission resources. We can adapt the deviation through a form of homeostatic plasticity, by increasing *β* when the deviation is below the channel capacity, and decreasing *β* when the deviation is above the channel capacity. Concretely, a simple update rule implements this idea:

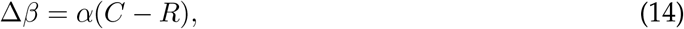

where *α* is a learning rate parameter. A similar adaptive gain modulation has been observed in neural circuits [36, 37, 38]. Mechanistically, this could be implemented by changes in background activity: when stimulus-driven excitation is high, the inhibition will also be high (the network is balanced), and the ensuing noise will effectively decrease the gain [39].

In the case where there are multiple stimuli, the same logic applies, but now we calculate the information rate over all the subpopulations of neurons (each coding a different stimulus). Specifically, the membrane potential becomes:

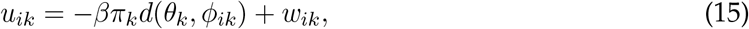

where *k* indexes both stimuli and separate subpopulations of neurons tuned to each stimulus location (or other stimulus feature that individuates the stimuli). As a consequence, *β* will tend to be smaller when more stimuli are encoded, because the same capacity constraint will be divided across more neurons.

### Memory maintenance

In delayed response tasks, the stimulus is presented transiently, and then probed after a delay. The channel thus needs to maintain stimulus information across the delay. Our model assumes that the membrane potential *u*_*i*_ maintains a trace of the stimulus across the delay. The persistence of this trace is determined by the gain parameter *β*. Because persistently high levels of stimulus-evoked activity may, according to Equation 13, increase the information rate above the channel capacity, the learning rule in Equation 14 will reduce *β* and thereby functionally decay the memory trace.

The circuit model does not commit to a particular mechanism for maintaining the stimulus trace. A number of suitable mechanisms have been proposed [40]. One prominent model posits that recurrent connections between stimulus-tuned neurons can implement an attractor network that maintains the stimulus trace as a bump of activity [41, 42]. Other models propose cell-intrinsic mechanisms [43, 44] or short-term synaptic modifications [45, 46]. All of these model classes are potentially compatible with the theory that population codes are optimizing a rate-distortion trade-off, provided that the dynamics of the memory trace conform to the equations given above. During time periods when no memory trace needs to be maintained, such as the intertrial interval (ITI) in delayed response tasks, we assume that the information rate is 0. Because the information rate is the *average* number of bits communicated across the channel, these “silent” periods effectively increase the achievable information rate during “active” periods (which we denote by *R*_*A*_). Specifically, if *T*_*A*_ is the active time (delay period length), and *T*_*S*_ is the silent time (ITI length), then the channel’s rate is given by:

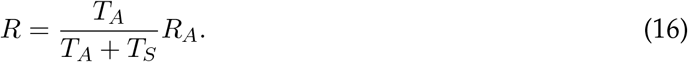

Equivalently, we can ignore the intervals in our model and simply rescale the channel capacity by (*T*_*A*_ + *T*_*S*_)*/T*_*A*_. This will allow us to model the effects of delay and ITI on performance in working memory tasks.

### Implications for working memory

#### Continuous report with circular stimuli

We apply the framework described above to the setting in which each stimulus is drawn from a circular space (e.g., color or orientation), *θ*_*k*_ ∈ (− *π, π*), which we discretize. Reconstruction errors are evaluted using a cosine distortion function:

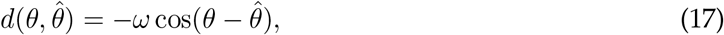

where *ω >* 0 is a scaling parameter. This implies that the input neurons have cosine tuning curves (Figure 1B). All of our subsequent simulations use the same tuning curves.

As an illustration of the model behavior in the continuous report task, we compare performance for set sizes 1 and 4. The optimal trade-off curves are shown in Figure 1C. For every point on the curve, the same information rate achieves a lower distortion for set size 1, due to the fact that all of the channel capacity can be devoted to a single stimulus (a hypothetical capacity limit is shown by the dashed horizontal line). In the circuit model, this higher performance is achieved by a narrow bump of population activity around the true stimulus (Figure 1D), compared to a broader bump when multiple stimuli are presented (Figure 1E).

#### Set size

One of the most fundamental findings in the visual working memory literature is that memory precision decreases with set size [47, 15, 48]. Our model asserts that this is the case because the capacity constraint of the system is divided across more neurons as the number of stimuli to be remembered increases, thus reducing the recall accuracy for any one stimulus. Figure 2A shows the distribution of recall error for different set sizes as published in previous work [15]. Figure 2D shows simulation results replicating these findings.

**Figure 2:**
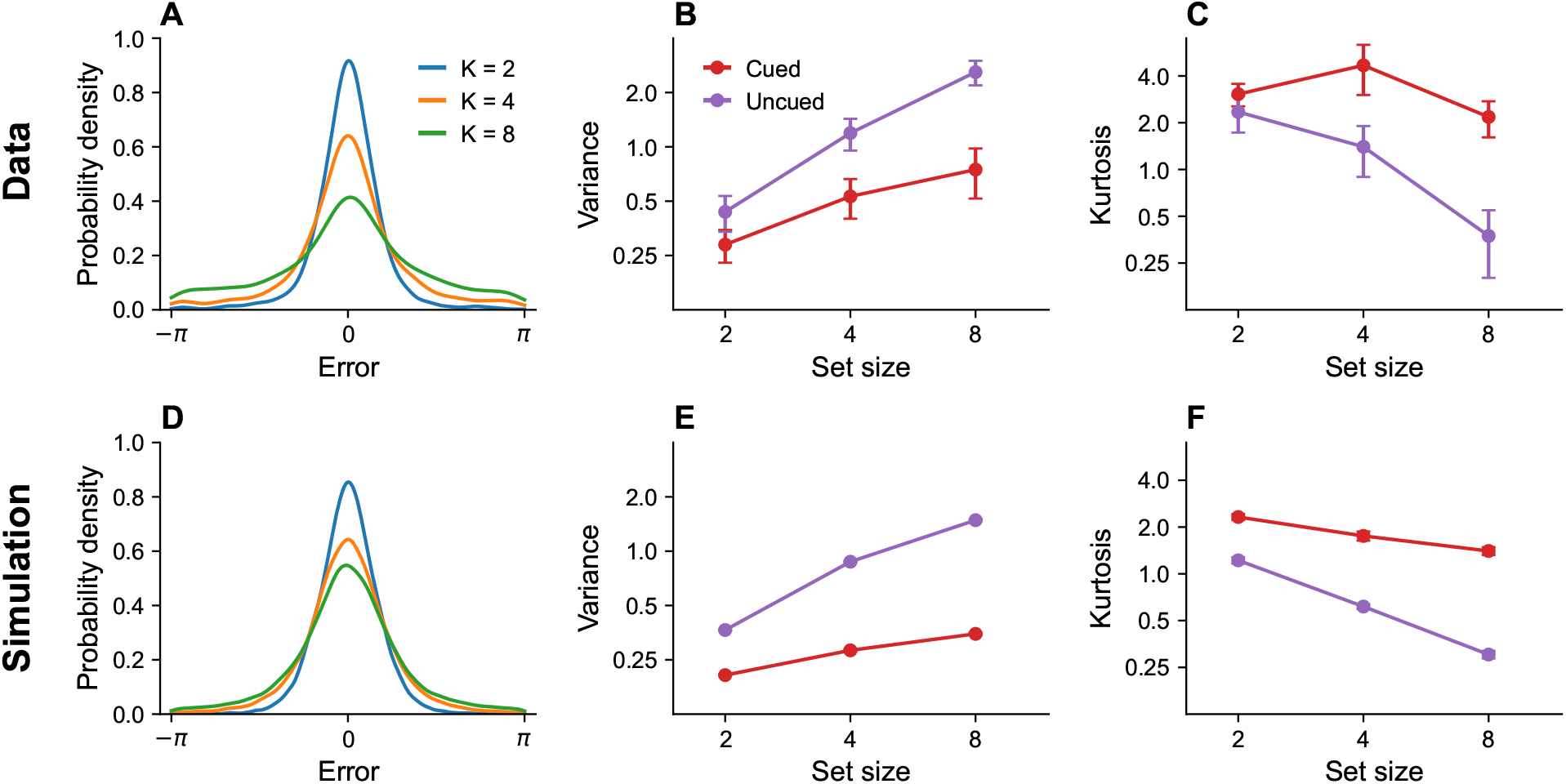
Set size effects and prioritization. (A) Error distributions for different set sizes, as reported in [15]. Error variability increases with set size. (B) Error variance as a function of set size for cued and uncued stimuli. Reports for cued stimuli have lower error variance. (C) Kurtosis as a function of set size for cued and uncued stimuli. (D, E, F) Simulation results replicating the observed effects. Error bars represent standard error of the mean.

#### Prioritization

Stimuli that are attentionally prioritized are recalled more accurately. For example, error variance is reduced by a cue that probabilistically predicts the location of the probed stimulus [15, 49]. In our model, the cue is encoded by the probing probability *π*_*k*_, which alters the expected distortion. This results in greater allocation of the capacity budget to cued stimuli than to uncued stimuli. Figure 2B and C show empirical findings, which are reproduced by our simulations shown in Figure 2E and F.

#### Timing

It is well-established that memory performance typically degrades with the retention interval [50, 51, 18, 52], although the causes of this degradation are controversial [53], and in some cases the effect is unreliable [54]. According to our model, this occurs because long retention intervals tax the information rate of the neural circuit. In order to stay within the channel capacity, the circuit reduces the gain parameter *β* for long retention intervals, thereby reducing the information rate and degrading memory performance.

Memory performance also depends on the intertrial interval, but in the opposite direction: longer intertrial intervals improve performance [26, 25]. The critical determinant of performance is in fact the ratio between the intertrial and retention intervals. Souza and Oberauer [26] found that performance in a color working memory task was similar when both intervals were short or both intervals were long. They also reported that a *longer* retention interval could produce *better* memory performance when it is paired with a longer intertrial interval. Figure 3 shows a simulation of the same experimental paradigm, reproducing the key results. This timescale invariance, which is also seen in studies of associative learning [55], arises as a direct consequence of Equation 16. Increasing the intertrial interval reduces the information rate, since no stimuli are being communicated during that time period, and can therefore compensate for longer retention intervals.

**Figure 3:**
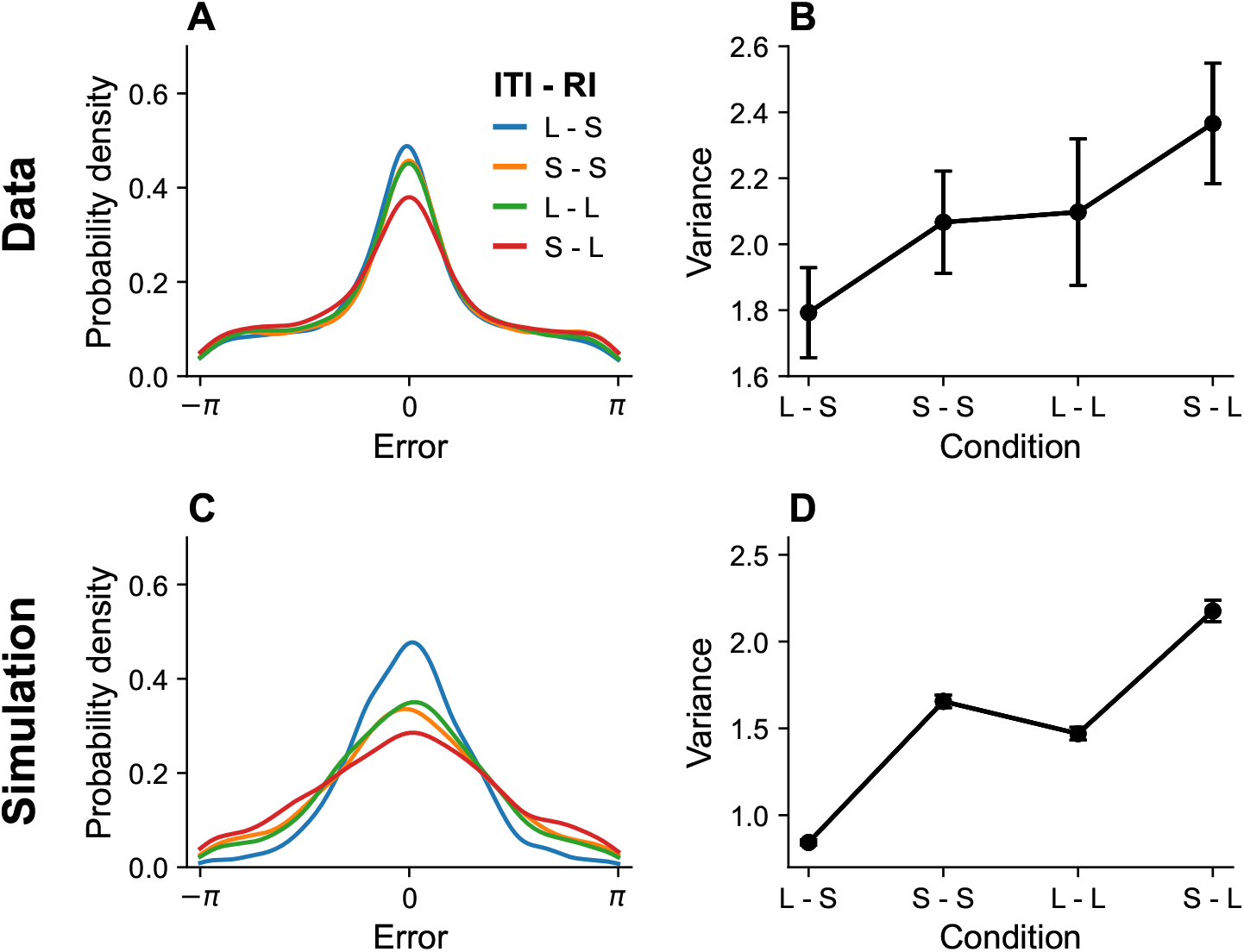
Timing effects. (A) Error distributions for different intertrial intervals (ITI) and retention intervals (RI), as reported in [26]. “S” denotes a short interval, and “L” denotes a long interval. Error variance as a function of timing parameters. Longer ITIs are associated with lower error variance, whereas longer RIs are associated with larger error variance. (C, D) Simulation results replicating the observed effects. Error bars represent standard error of the mean.

#### Serial dependence

Working memory recall is biased by recent stimuli, a phenomenon known as *serial dependence* [58, 59, 27, 60]. Recall is generally attracted toward recent stimuli, though some studies have reported repulsive effects when the most recent and current stimulus differ by a large amount [61, 27]. Our theory explains serial dependence as a consequence of the marginal firing rate of the output cells, which biases the excitatory input *u*_*i*_ (see Equation 10). Because the marginal firing rate is updated incrementally, it will reflect recent stimulus history.

An important benchmark for theories of serial dependence is the finding that it increases with the retention interval and decreases with intertrial interval [27]. These twin dependencies are reproduced by our model (Figure 4). Our explanation of serial dependence is closely related to our explanation of timing effects on recall error: the strength of serial dependence varies inversely with the information rate, which in turn increases with the intertrial interval and decreases with the retention interval. Mechanistically, this effect is mediated by adjustments of the gain parameter *β* in order to keep the information rate near the channel capacity.

**Figure 4:**
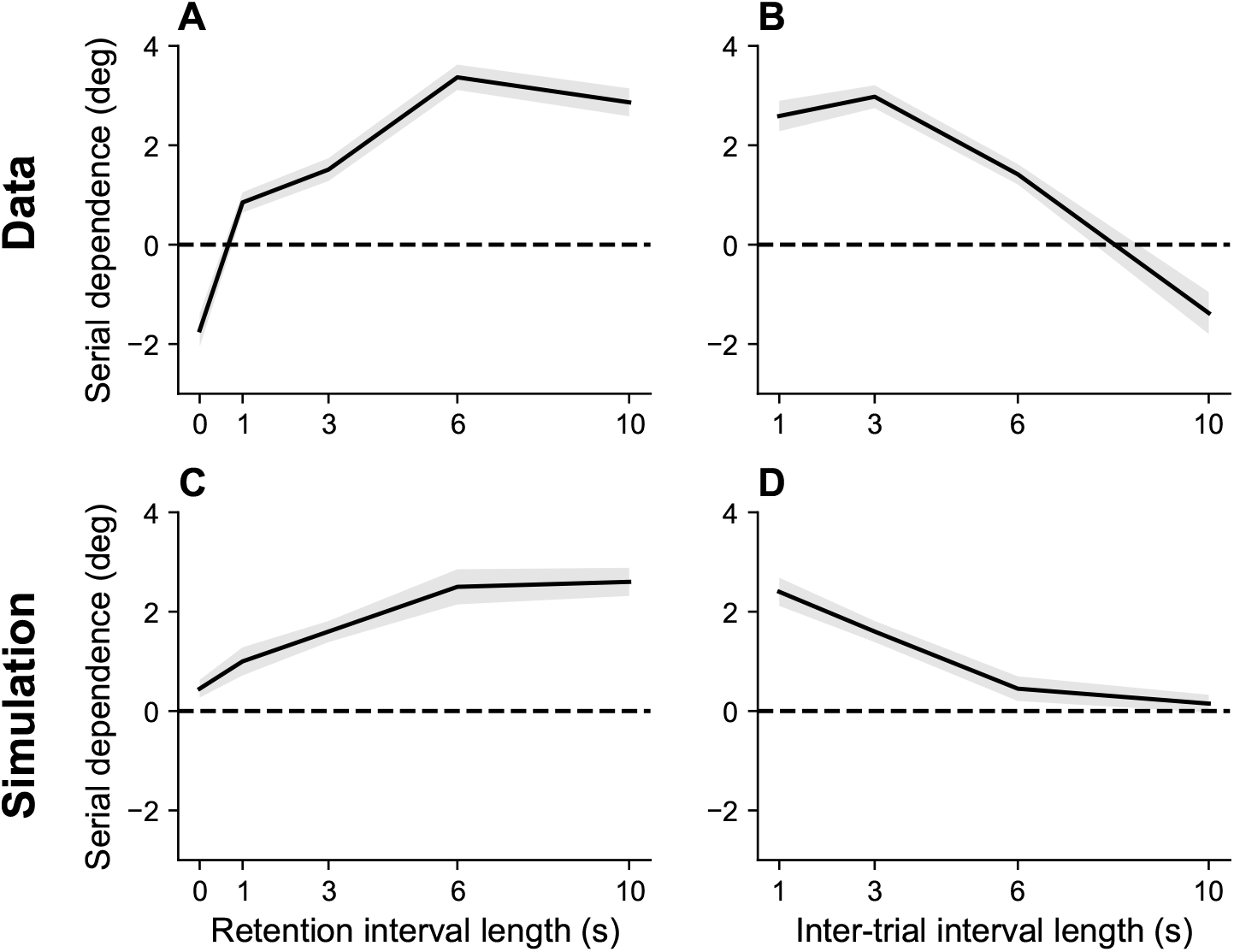
Serial dependence as a function of retention interval and intertrial interval. (A) Serial dependence increases with the retention interval until eventually reaching an asymptote, as reported in [27]. Serial dependence is quantified as the peak-to-peak amplitude of a derivative of Gaussian (DoG) tuning function fitted to the data using least squares (see Methods). (B) Serial dependence decreases with intertrial interval. (C,D) Simulation results. Shaded area corresponds to standard error of the mean.

Serial dependence has also been shown to build up over the course of an experimental session [56]. This is hard to explain in terms of theories based on purely short-term effects, but it is consistent with our account in terms of the bias induced by the marginal firing rate. Because this bias reflects continuous incremental adjustments, it integrates over the entire stimulus history, thereby building up over the course of an experimental session.

If, as we hypothesize, serial dependence reflects a capacity limit, then we should expect it to increase with set size, since *β* must decrease to stay within the capacity limit. To the best of our knowledge, this prediction has not been tested. We confirmed this prediction for color working memory using a large dataset reported in [51]. Figure 6 shows that the attractive bias for similar stimuli on consecutive trials is stronger when the set size is larger (*p <* 0.05, group permutation test).

**Figure 5:**
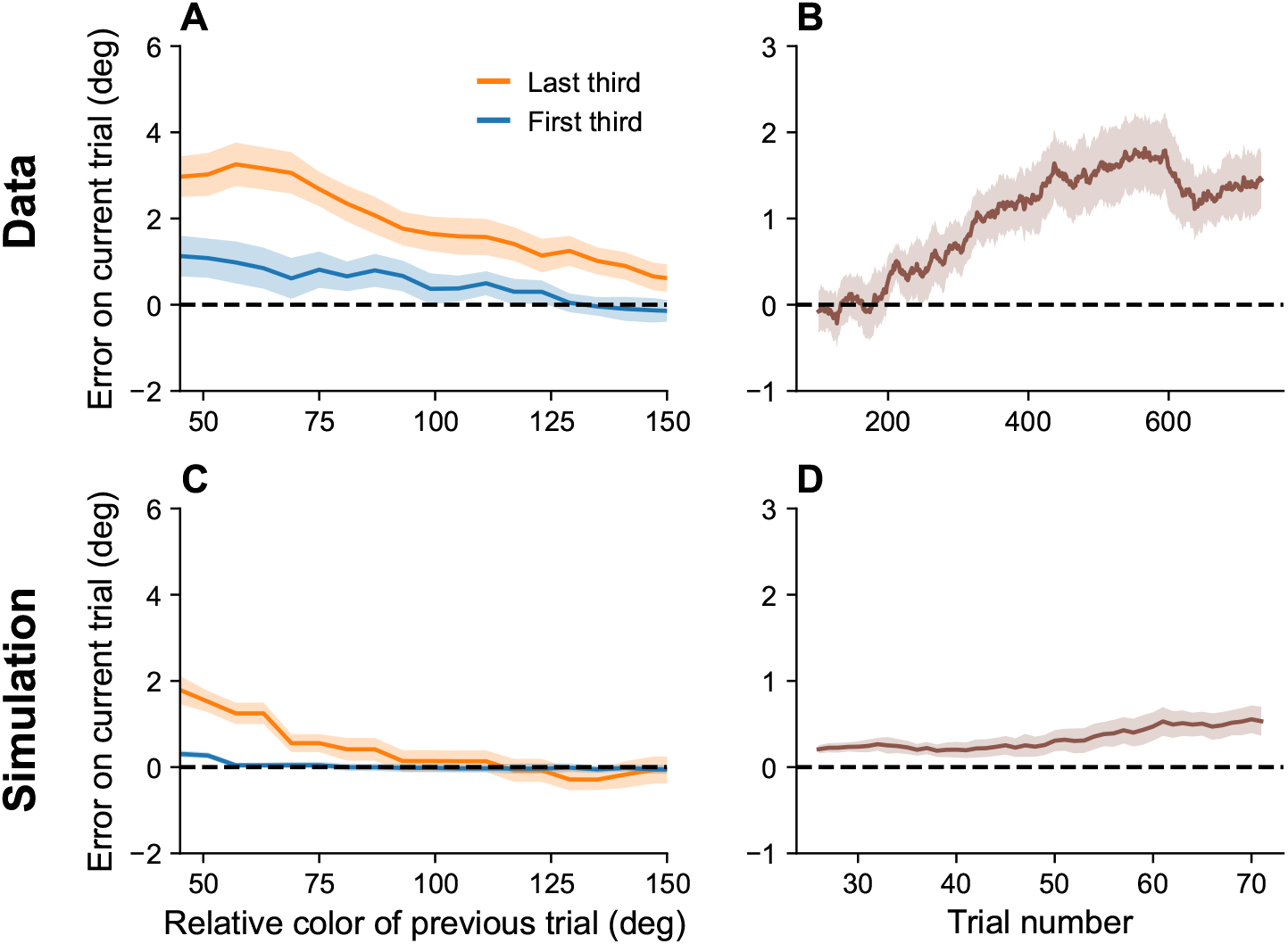
Serial dependence builds up during an experiment. (A) Serial dependence computed using first third (blue) and last third (orange) of the trials within a session, as reported in [56]. Data shown here were originally reported in [57]. To obtain a trial-by-trial measure of serial dependence, we calculated the folded error as described in [56] (see Methods). Positive values indicate attraction to the last stimulus, while negative values indicate repulsion. Serial dependence is stronger in the last third of the trials in the experiment compared to the first third. (B) Serial dependence increases over the course of the experimental session, computed here with a sliding window of 200 trials. (C, D) Simulation results. Shaded area corresponds to standard error of the mean.

**Figure 6:**
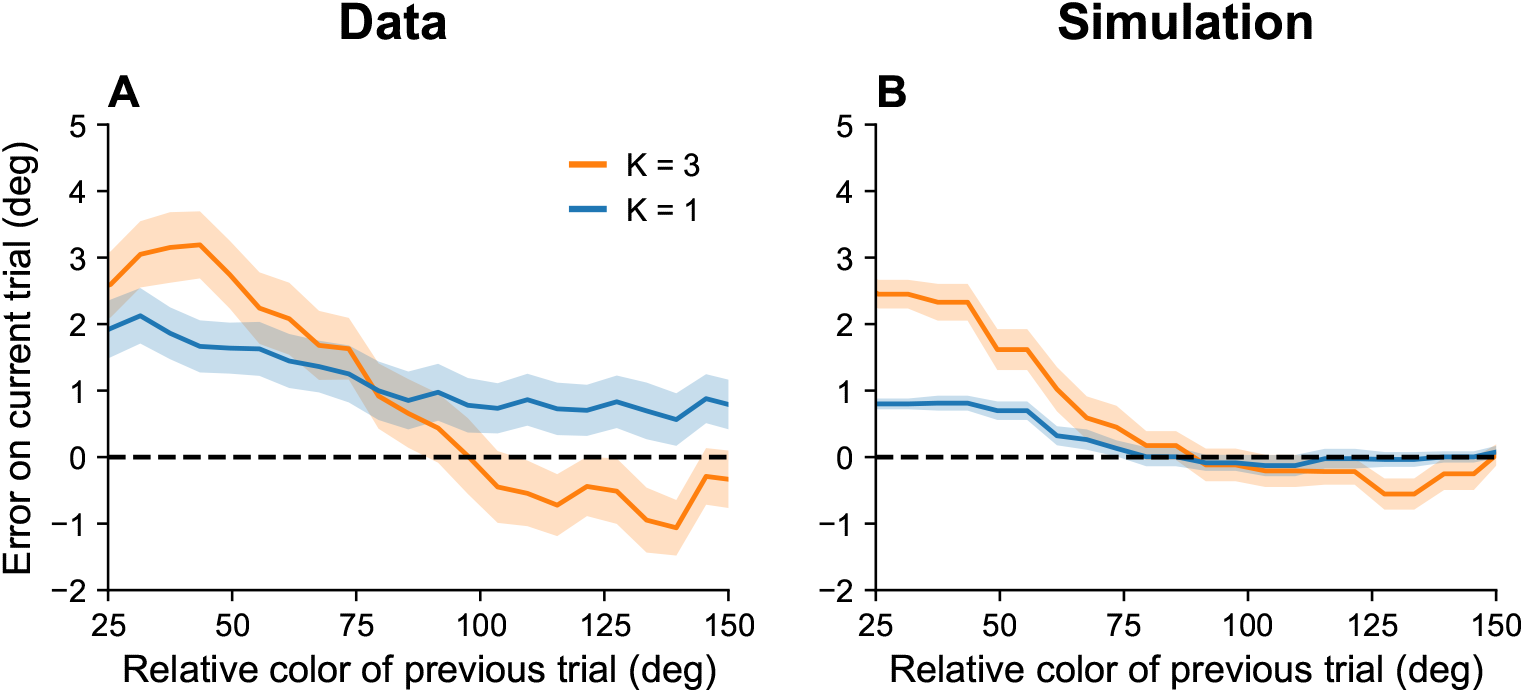
Serial dependence increases with set size. (A) Serial dependence (quantified using folded error) for set sizes *K* = 1 (blue) and *K* = 3 (orange), using data originally reported in [51]. Serial dependence computed as the peak amplitude of a DoG tuning function fitted to the data using least squares is stronger for larger set sizes (see Methods). (B) Simulation results. Shaded area corresponds to standard error of the mean.

#### Systematic biases

Working memory exhibits systematic biases towards stimuli that are shown more frequently than others [51]. Moreover, these biases increase with the retention interval, and build up over the course of an experimental session. Our interpretation of serial dependence, which also builds up over the course of a session, suggests that these two phenomena may be linked (see also [62]).

Our theory posits that, over the course of the experiment, the marginal firing rate asymptotically approaches the distribution of presented stimuli (assuming there are no inhomogeneities in the distortion function). Thus, the neurons corresponding to high-frequency stimuli become more excitable than others and bias recall towards their preferred stimuli. This bias is amplified by lower effective capacities brought about by longer retention intervals. Figure 7 shows simulation results replicating these effects.

**Figure 7:**
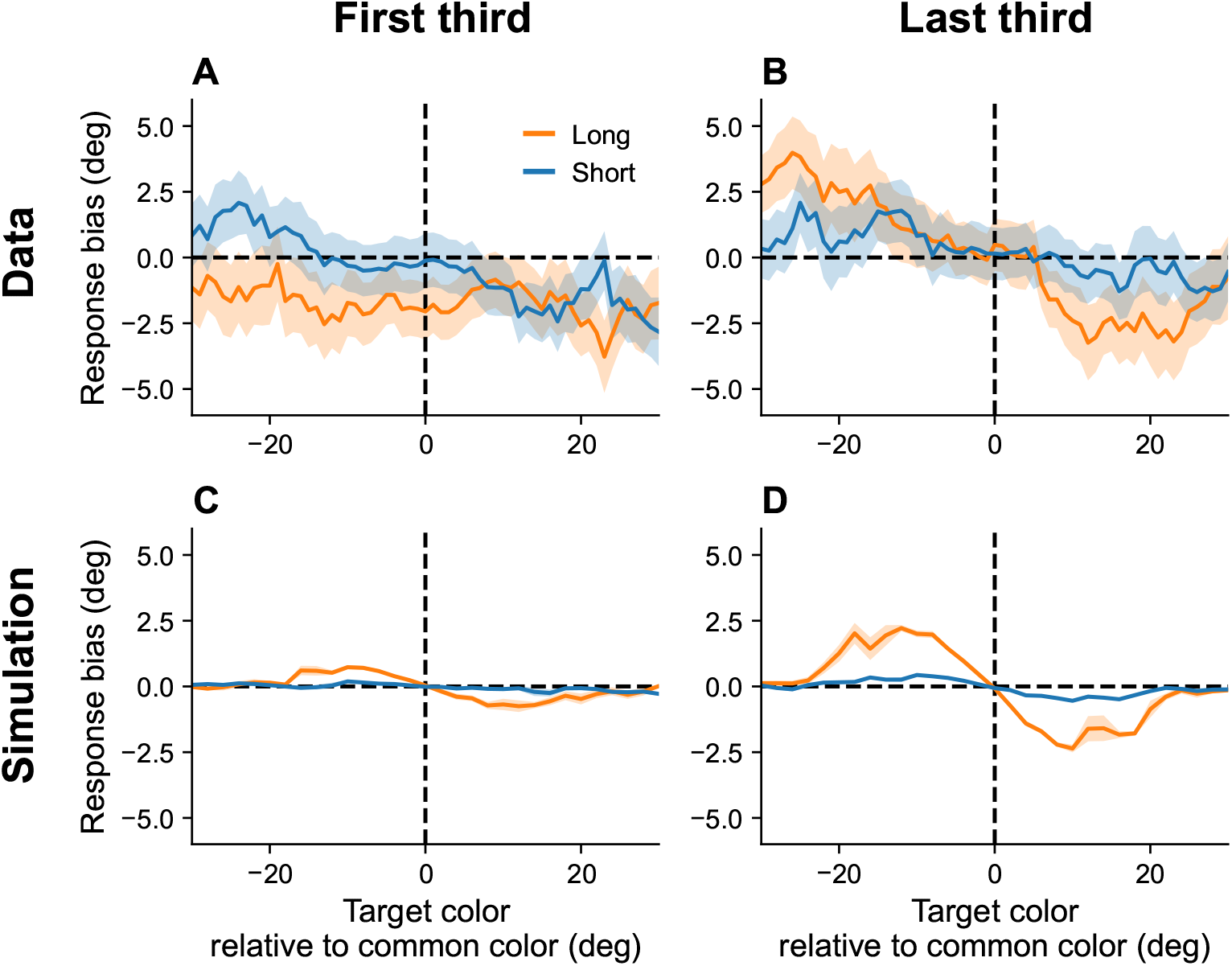
Continuous reports are biased towards high frequency colors. (A, B) Bias for targets around common colors during the first (Panel A) and last (Panel B) third of the session, as reported in [51]. Bias refers to the difference between the stimulus and the mean reported color. X axis is centered around high-frequency colors. Bias increases with RI length (blue = short RI, orange = long RI). Bias also increases as the experiment progresses. (C, D) Simulation results. Shaded area corresponds to standard error of the mean.

#### Variations in gain

Equation 14 predicts that operating below the channel capacity will lead to an increase in the gain term *β*, which, in turn, leads to a higher information rate and better memory performance. Therefore, our model predicts that recall accuracy should improve after a period of poor memory performance, and degrade after a period of good memory performance. At the neural level, the model predicts that error will tend to be lower when gain (*β*) is higher.

We tested these predictions by re-analyzing the monkey neural and behavioral data reported in [61] (*N* = 2). Squared error was significantly lower following higher-than-average error than following lower-than-average error (linear mixed model, *p <* 0.001; Figure 8A).

**Figure 8:**
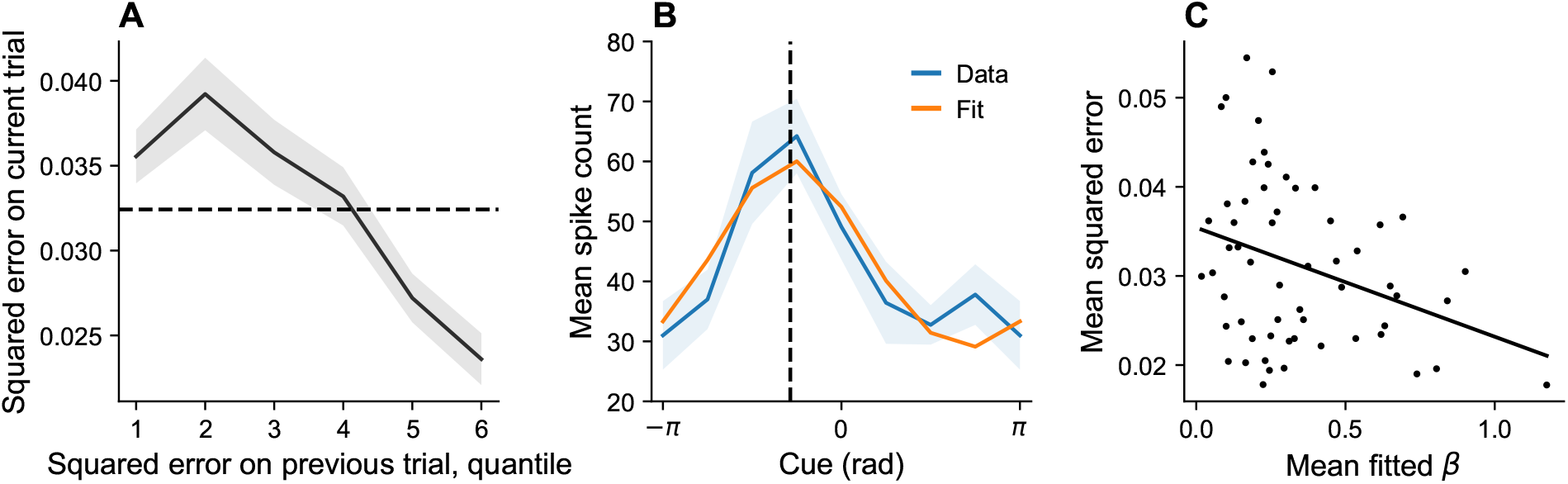
Dynamic variation in memory precision and neural gain. (A) Mean squared error on current trial, classified by quantiles of squared error on previous trial. Squared error tends to be above average (dashed black line) following low squared error on the previous trial, and tends to be below average following large squared error on the previous trial. (B) Orientation tuning curve (orange) fitted to mean spike count (blue) during the retention interval, shown for one example neuron. The neuron’s preferred stimulus (dashed black line) corresponds to the peak of the tuning curve. Shaded region corresponds to standard error of the mean. (C) Mean squared error for different sessions plotted against mean fitted *β*. According to our theory, *β* plays the role of a gain control on the stimulus. Consistent with this hypothesis, memory error decreases with *β*.

In order to estimate the neural gain, we first inferred the preferred stimulus of each neuron by fitting a bell-shaped tuning function to its spiking behavior (Equation 21, Figure 8B). We then performed Poisson regression to fit a *β* for each neuron (Equation 22). Model comparison using the Bayesian information criterion (BIC) established that both the distortion function (which captures driving input) and spiking history were significant predictors of spiking behavior (full model: 54, 545; no history: 59, 163; neither distortion nor history: 67, 903). We then examined the relationship between neural gain and memory precision across sessions, finding that session-specific mean squared error was negatively correlated with the average *β* estimate (*r* = −0.32, *p <* 0.02; Figure 8C).

## Discussion

We have shown that a simple population coding model with spiking neurons can solve the channel design problem: signals passed through the spiking network are transmitted with close to the minimum achievable distortion under the network’s capacity limit. We focused on applying this general model to the domain of working memory, unifying several seemingly disparate aspects of working memory performance: set size effects, stimulus prioritization, serial dependence, approximate timescale invariance, and systematic bias. Our approach builds a bridge between biologically plausible population coding and prior applications of rate-distortion theory to human memory [1, 2, 9, 4, 5, 10].

### Relationship to other models

The hypothesis that neural systems are designed to optimize a rate-distortion trade-off has been previously studied through the lens of the information bottleneck method [63, 64, 65, 66], a special case of rate-distortion theory in which the distortion function is derived from a compression principle. Specifically, the distortion function is defined as the Kullback-Leibler divergence between *P* (*θ*^′^| *θ*) and 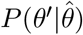, where *θ*^′^ denotes the probed stimulus. This distortion function applies a “soft” penalty to errors based on how much probability mass the channel places on each stimulus. The expected distortion is equal to the mutual information between *θ*^′^ and 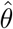. Thus, the information bottleneck method seeks a channel that maps the input *θ* into a compressed representation 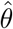 satisfying the capacity limit, while preserving information necessary to predict the probe *θ*^′^.

As pointed out by Leibfried and Braun [67], using the Kullback-Leibler divergence as the distortion function leads to a harder optimization compared to classical rate-distortion theory because 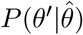 depends on the channel distribution, which is the thing being optimized. One consequence of this dependency is that minimizing the rate-distortion objective using alternating optimization (in the style of the Blahut-Arimoto algorithm) is not guaranteed to find the globally optimal channel. It is possible to break the dependency by replacing 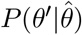 with a reference distribution that does not depend on the channel. This turns out to strictly generalize rate-distortion theory, because an arbitrary choice of the reference distribution allows one to recover any lower-bounded distortion function up to a constant offset [67]. However, existing spiking neuron implementations of the information bottleneck mthod [64, 65] do not make use of such a reference distribution, and hence do not attain the same level of generality.

Leibfried and Braun [67] propose a spiking neuron model that explicitly optimizes the rate-distortion objective function for arbitrary distortion functions. Their approach differs from ours in several ways. First, they model a single neuron, rather than a population. Second, they posit that the channel optimization is realized through synaptic plasticity, in contrast to the intrinsic plasticity rule that we study here. Third, they treat the gain parameter *β* as fixed, whereas we propose an algorithm for optimizing *β*.

### Open questions

A cornerstone of our approach is the assumption that the neural circuit responsible for working memory dynamically modifies its output to stay within a capacity limit. What, at a biological level, is the nature of this capacity limit? Spiking activity accounts for a large fraction of cortical energy expenditure [68, 69]. Thus, a limit on the overall firing rate of a neural population is a natural transmission bottleneck. Previous work on energy-efficient coding has similarly used the cost of spiking as a constraint [34, 70, 71]. One subtlety is that the capacity limit in our framework is an upper bound on the stimulus-driven firing rate *relative* to the average firing rate (on a log scale). This means that the average firing rate can be high provided the stimulus-evoked transients are small, consistent with the observation that firing rate tends to be maintained around a set point rather than minimized [36, 37, 38]. The set point should correspond to the capacity limit.

The next question is how a neural circuit can control its sensitivity to inputs in such a way that the information rate is maintained around the capacity limit. At the single neuron level, this might be realized by adaptation of voltage conductances [70]. At the population level, neuromodulators could act as a global gain control. Catecholamines (e.g., dopamine and norepinephrine), in particular, have been thought to play this role [72, 73]. Directly relevant to this hypothesis are experiments showing that local injection of dopamine receptor antagonists into the prefrontal cortex impaired performance in an oculomotor delayed response task [74].

Our model can be extended in several ways. One, as already mentioned, is to develop a biologically plausible implementation of gain adaptation, either through intrinsic or neuromodulatory mechanisms. A second direction is to consider channels that transmit a compressed representation of the input. Previous work has suggested that working memory representations are efficient codes that encode some stimuli with higher precision than others [75, 20]. Finally, an important direction is to enable the model to handle more complex memoranda, such as natural images. Recent applications of large-scale neural networks, such as the variational autoencoder, to modeling human memory hold promise [5, 76], though linking these to more realistic neural circuits remains a challenge.

## Methods

We re-analyzed 6 datasets of monkey and human subjects performing a delayed response task. The detailed experimental procedures can be found in the original reports [15, 26, 61, 56, 51, 46]. In 3 of the 6 datasets, one or multiple colors were presented on a screen at equally spaced locations. After a retention interval, during which the cues were no longer visible, subjects had to report the color at a particular cued location, measured as angles on a color wheel. In one dataset, angled color bars were presented, and the angle of the bar associated with a cued color had to be reported [15]. In the two last datasets, only the location of a black cue on a circle had to be remembered and reported [61, 46].

### Set size and stimulus prioritization

Human subjects (*N* = 7) were presented with 2, 4 or 8 color stimuli at the same time. On each trial, one of the locations was cued before the appearance of the stimuli. Cued locations were 3 times as likely to be probed [15].

We computed trial-wise error as the circular distance between the reported angle and the target angle, separately for each set size and cuing condition. We then calculated circular variance and kurtosis as presented in the original paper, using the following equations:

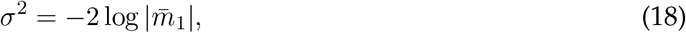

and

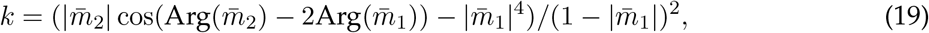

where 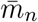 is the *n*th uncentered trigonometric moment.

### Timing effects

Human subjects (*N* = 36) were presented with 6 simultaneous color stimuli and had to report the color at a probed location as an angle on a color wheel. The RI and ITI lengths varied across sessions (RI: 1 or 3 seconds, ITI: 1 or 7.5 seconds) [26].

### Serial dependence increases with retention interval and decreases with intertrial interval

Human subjects (*N* = 55) were presented with a black square at a random position on a circle and had to report the location of the cue [46]. The RI and ITI were varied across blocks of trials (RI: 0, 1, 3, 6, or 10 seconds, ITI: 1, 3, 6 or 10 seconds). For each block and subject, we computed serial dependence as the peak-to-peak amplitude of a derivative of Gaussian (DoG) function fit to the data. The DoG function is defined as follows:

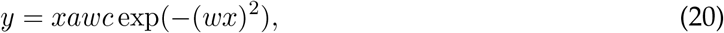

where *y* is the trial-wise error, *x* is the relative circular distance to the target angle of the previo<u>u</u>s trial, *a* is the amplitude of the DoG peak, *w* is the width of the curve, and *c* is the constant 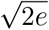, chosen such that the peak-to-peak amplitude of the DoG fit—the measure of serial dependence in [46]—is exactly 2*a*.

### Build-up of serial dependence

Human subjects (*N* = 12) performed a delayed continuous report task with one item [57]. Following [56], we obtained a trial-by-trial measure of serial dependence using their definition of folded error.

Let *θ*_*d*_ denote the circular distance between the angle reported on the previous trial and the target angle on the current trial. In order to aggregate trials with negative *θ*_*d*_ (preceding target is located clockwise to current target) and trials with positive *θ*_*d*_ (preceding target is located counter-clockwise to current target), we computed the folded error as 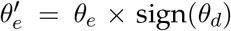, where *θ*_*e*_ is the circular distance between the reported angle and the target angle. Positive 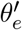 corresponds to attraction to the previous stimulus, whereas negative 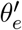 corresponds to repulsion.

We excluded trials with absolute errors larger than *π/*4. We then computed serial bias as the average folded error in sliding windows of width *π/*2 rad and steps of *π/*30 rad. We repeated this procedure separately for the trials contained in the first and last third of all sessions. Finally, we computed the increase in serial dependence over the course of a session using a sliding window of 200 trials on the folded error.

### Serial dependence increases with set size

We re-analyzed the dataset collected by [51], experiment 1a, in which human subjects (*N* = 90) performed a delayed response task with 1 or 3 items.

We calculated folded error using the procedure mentioned above. We excluded trials with absolute errors larger than *π/*4. We then computed serial bias as the average folded error in sliding windows of width *π/*4 rad and steps of *π/*30 rad. We repeated this procedure separately for the trials with *K* = 1 or *K* = 3 items. In order to test whether serial dependence was stronger for one of the set size conditions, we performed a permutation test: We shuffled the entire dataset and partitioned it into two groups of size *S*_*K*=1_ and *S*_*K*=3_, where *S*_*K*=*k*_ denotes the number of trials recorded for the set size condition *K* = *k*. We fitted a DoG curve (Equation 20) to each partition using least squares and computed the difference between the peak amplitude of the two fits. We repeated this process 20, 000 times. We then calculated the p-value as the proportion of shuffles for which the difference between the peak amplitudes was equal to or larger than the one computed using the unshuffled dataset.

### Continuous reports are biased towards high frequency colors

Human subjects (*N* = 120) performed a delayed continuous report task with a set size of 2 [51]. On each trial, the RI was either 0.5 or 4 seconds. The stimuli were either drawn from a uniform distribution or from a set of 4 equally-spaced bumps of width *π/*9 rad with equal probability. The centers of each bump were held constant for each subject.

We defined systematic bias as mean error versus distance to the closest bump center and computed it in sliding windows of width *π/*45 rad and steps of *π/*90 rad, as done in the original study. We repeated this procedure separately for the trials with *RI* = 0.5*s* or *RI* = 4*s*, and for the first and last third of trials within a session.

### Simulation parameters

We used the following parameters for all simulations, unless stated otherwise: *N* = 1000, *K* = 1, *C* = 0.9, *ω* = 1, *η* = 3 ×10^−6^, *c* = 0.5, *α* = 10^−3^,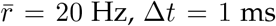. Spikes contributed to intrinsic synaptic plasticity for 10 timesteps. Weights *w* were clipped to be in the range [−12, 0]. *β* was initialized at *β*_0_ = 20 and clipped to be in the range [0, 200]. *C* was the only free parameter we allowed to vary across simulations, and we chose *C* = 0.1 to fit the data in [26].

In order to account for the higher probing probability of the cued stimulus in [15], we used 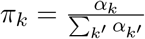 with *α*_priority_ = {1.5_*K*=2_, 2.2_*K*=4_, 3.0_*K*=8_} and *α*_*k*_ = 1 otherwise, as reported in [15].

### Dynamics of memory precision and neural gain

We re-analyzed the behavioral and neural dataset collected in [61]. Since neural recordings were not available for all trials within a session, we ignored sessions in which only a subset of the 8 potential cues were displayed.

We sorted the squared error on trial *t* (denoted by 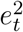) based on 6 quantiles of the squared error on the previous trial. We then defined the indicator variable 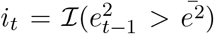, taking the value +1 if the squared error on the previous trial was larger than the mean squared error, and −1 otherwise. We then fit the linear mixed model 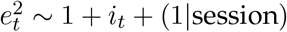.

In order to infer the preferred stimulus of each recorded neuron, we used a least squares approach to fit the mean spike count for each presented stimulus and neuron to a bell-shaped tuning function:

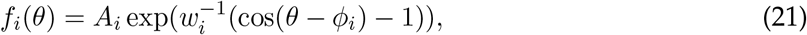

where *θ* is the presented stimulus, *A*_*i*_ and *w*_*i*_ control the amplitude and width of the tuning function, respectively, and *ϕ*_*i*_ is the preferred stimulus of neuron *i* [15].

We then fitted the neural data by performing Poisson regression for each neuron using the following model:

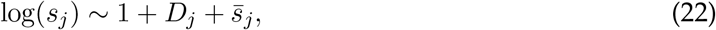

where *s*_*j*_ is the number of spikes emitted by the neuron on trial *j, D*_*j*_ is the expected distortion between the stimulus *θ*_*j*_ and the neuron’s preferred stimulus, and 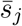 is an exponential moving average of the neuron’s spike history with decay rate 0.8. We discarded 3 neurons for which the fitted *β* was negative and 1 neuron for which the fitted *β* was larger than 5 standard deviations above the mean of the fitted values. In order to ascertain the utility of the different regressors, we fitted another model without the history term, and another without both the distortion and history terms, and compared them based on their Bayesian Information Criterion (BIC).

### Source code

All simulations and analyses were performed using Julia, version 1.6.2. Source code can be found at https://github.com/amvjakob/wm-rate-distortion.

## Acknowledgments

Johannes Bill and Chris Bates generously provided constructive feedback and discussion. We are also grateful to Dan Bliss and Matt Panichello for sharing data. This research was supported by a Bertarelli Fellowship and by the Center for Brains, Minds, and Machines (funded by NSF STC award CCF-1231216).

